# NEIL2 plays a critical role in limiting inflammation and preserving genomic integrity in *H. pylori*-infected gastric epithelial cells

**DOI:** 10.1101/687962

**Authors:** Ayse Z Sahan, Tatiana Venkova, Ibrahim M. Sayed, Ellen J Beswick, Victor E. Reyes, Irina Pinchuk, Debashis Sahoo, Pradipta Ghosh, Tapas K Hazra, Soumita Das

## Abstract

The accumulation of *Helicobacter pylori* infection-induced oxidative DNA damage in gastric epithelial cells is a risk factor for developing gastric cancer (GC); however, the underlying mechanisms remain poorly understood. Here we report that the suppression of NEIL2, an oxidized base-specific mammalian DNA glycosylase, is one such mechanism via which *H. pylori* infection may fuel the accumulation of DNA damage during the initiation and progression of GC. Using a combination of cultured cell lines and primary cells, we show that expression of NEIL2 is significantly down-regulated after *H. pylori* infection; such down-regulation was also seen in human gastric biopsies. The *H. pylori* infection-induced down-regulation of NEIL2 is specific, as *Campylobacter jejuni* has no such effect. Using gastric organoids isolated from the murine stomach in co-culture studies with live bacteria mimicking the infected stomach lining, we found that *H. pylori* infection was associated with IL-8 production; this response was more pronounced in *Neil2* knockout (KO) mouse cells compared to wild type (WT) cells, suggesting that NEIL2 suppresses inflammation under physiological conditions. Interestingly, DNA damage was significantly higher in *Neil2* KO mice compared to WT mice. *H. pylori*-infected *Neil2* KO mice showed higher inflammation and more epithelial cell damage. Computational analysis of gene expression profiles of repair genes in gastric specimens showed the reduction of *Neil2* level is linked to the GC progression. Taken together, our data suggest that down-regulation of NEIL2 is a plausible mechanism by which *H. pylori* infection derails DNA damage repair, amplifies the inflammatory response and initiates GCs.

## Introduction

The World Health Organization and the American Cancer Society (ACS) estimate that Gastric Cancer (GC) is the fourth most prevalent cancer worldwide, and the third leading cause of death in the United States (1). Recognized risk factors for developing GCs include genetic predisposition (variants in DNA repair genes have been reported as key factors in GC susceptibility and survival) (2,3), dietary factors (for example, high salt intake), obesity, smoking, and *Helicobacter pylori (H. pylori)* infection (4,5). *H. pylori* is a gram-negative, microaerophilic bacterium that infects the gastric mucosa, causing gastritis and often leading to devastating complications such as peptic ulcers and GC. *H. pylori* infection increases the risk for developing GCs by 5-6 fold (6). Treatment and eventual eradication of the infection could reduce the risk of development of these complications, but this cannot completely erase the possible threat. Additionally, an increasing number of reports of antibiotic resistance among *H. pylori* strains. This makes complete eradication of *H. pylori* infection challenging and recurrent infections are common without extreme or noticeable symptoms (5,7,8).

As for the pathogenesis of GCs, infection with *H. pylori* of both gastric cell lines and primary gastric epithelial cells has been shown to trigger a unique DNA damage pattern, one that preferentially targets transcribed regions and regions proximal to telomeres (6,9). However, how *H. pylori* infection results in the accumulation of DNA damage remains unknown. Generally, the bacterial infection-induced inflammation leads to the generation of reactive oxygen species (ROS) that in turn cause oxidative genome damage in the host cells. Without timely repair, the mutagenic DNA lesions can cause several pathologies, including cancer (10-13). Oxidized DNA bases are primarily repaired via the DNA Base Excision Repair (BER) pathway (14,15). BER is a multistep process; it is initiated with recognition and excision of the lesion by a family of repair proteins called DNA glycosylases (DGs). All four DNA bases are subjected to oxidative modifications leading to approximately two dozen lesions, and only 5 oxidized base-specific DNA glycosylases with overlapping substrate specificities have been characterized so far in mammalian cells. Among the five, OGG1 and NTH1, identified and characterized earlier, primarily remove oxidized purines and pyrimidines respectively and were thought to be the major mammalian DNA glycosylases. However, null mice of these two DGs did not show any visible phenotype (14,16-22). Several other groups and we then found that mammalian cells possess a new family of DNA glycosylases, and named them *E. coli* Nei-like (NEIL1-3). NEILs are capable of removing both purines and pyrimidines; however, these DGs are unique with respect to DNA damage recognition, as OGG1 and NTH1 remove the damage from duplex DNA, whereas NEILs prefer to initiate excision of lesions that are located in a single-stranded region, a replication fork or transcription bubble mimic (13,23,24). Indeed, it was found that NEIL1 and NEIL3 are mostly involved in repairing the damage in replicating cells, and NEIL2 primarily removes the oxidized DNA bases from the transcribed regions (24). Recently, copy number variation analysis of the COSMIC and TCGA databases has clearly shown that, among all DNA repair proteins, NEIL2 is the most affected in various cancers, and its level is generally low (25). A mutant or functional variant of NEIL2 was also reported as a risk factor for lung cancer and squamous cell carcinoma in the oral cavity (26-29), and the loss of NEIL2 is associated with poor survival in patients with estrogen-receptor positive breast cancer (30).

Several reports indicate that *H pylori*-induced ROS can cause DNA damage leading to mutation (31,32). Hence, it is expected that aberrant expression and/or compromised activities of the DGs will be contributing factors for various pathologies, including carcinogenesis. Surprisingly, despite the accumulation of significantly higher amounts of mutagenic 8-oxoG in Ogg1-/-mice, these animals did not develop tumors when exposed to KBrO3, an oxygen radical forming agent (33). Moreover, Ogg1-null mice are resistant to *H. pylori*-induced inflammation. These data suggest that oxidized DNA base lesions alone are not sufficient; some other factor(s) would be necessary for developing tumors in the affected tissues.

Based on the link between NEIL2 and various types of cancers, we hypothesized that *H. pylori* infection may trigger the accumulation of mutations via dysregulation of NEIL2. Using the stem cell-based *ex vivo* technique, in which gastric organoids were isolated from the stomach of WT and *Neil2*-KO mice and an *in vivo* infection model, we report here that the level of NEIL2s, but not that of other DGs, is significantly decreased upon *H. pylori* infection, leading to increased DNA damage accumulation and inflammatory responses. Because ROS, DNA damage, and inflammation are closely intertwined, and because NEIL2 is an anti-inflammatory protein in addition to its role in DNA repair, we conclude that the combinatorial effect of accumulation of oxidized DNA lesions along with chronic inflammation likely plays a critical role in the initiation and progression of *H pylori*-associated GCs.

## Results

### H. pylori infection is associated with altered expression of NEIL2 in a dose and time-dependent manner

To determine the status of BER proteins in gastric epithelial cells after *H. pylori* infection, AGS cells were transfected with the FLAG-tagged expression plasmid of OGG1, NEIL1, NEIL2, and NTH1 followed by 24 h of infection with *H. pylori*. Surprisingly, NEIL2’s level, but not that of other DNA glycosylases, was significantly decreased in the AGS cell lysate (Fig. 1A). Moreover, endogenous expression of NEIL2 in AGS cells after *H. pylori* infection was also significantly decreased compared to the OGG1 (Fig. 1B). Furthermore, the expression of NEIL2 following *H. pylori* infection was decreased in a time-dependent manner. The expression started to decrease after 6 h post infection and became undetectable at 24 and 48 h (Fig. 1C). To test the effect of *H. pylori* inoculum on NEIL2 expression, AGS cells were infected with *H. pylori* at MOI of 10 and 100, and the expression of NEIL2 was determined by Western blot. The reduction of NEIL2 expression was more pronounced in AGS lysates at higher bacterial load (Fig. 1D), indicating that the association of *H. pylori* infection is linked to down-regulation of NEIL2.

**Figure 1.**
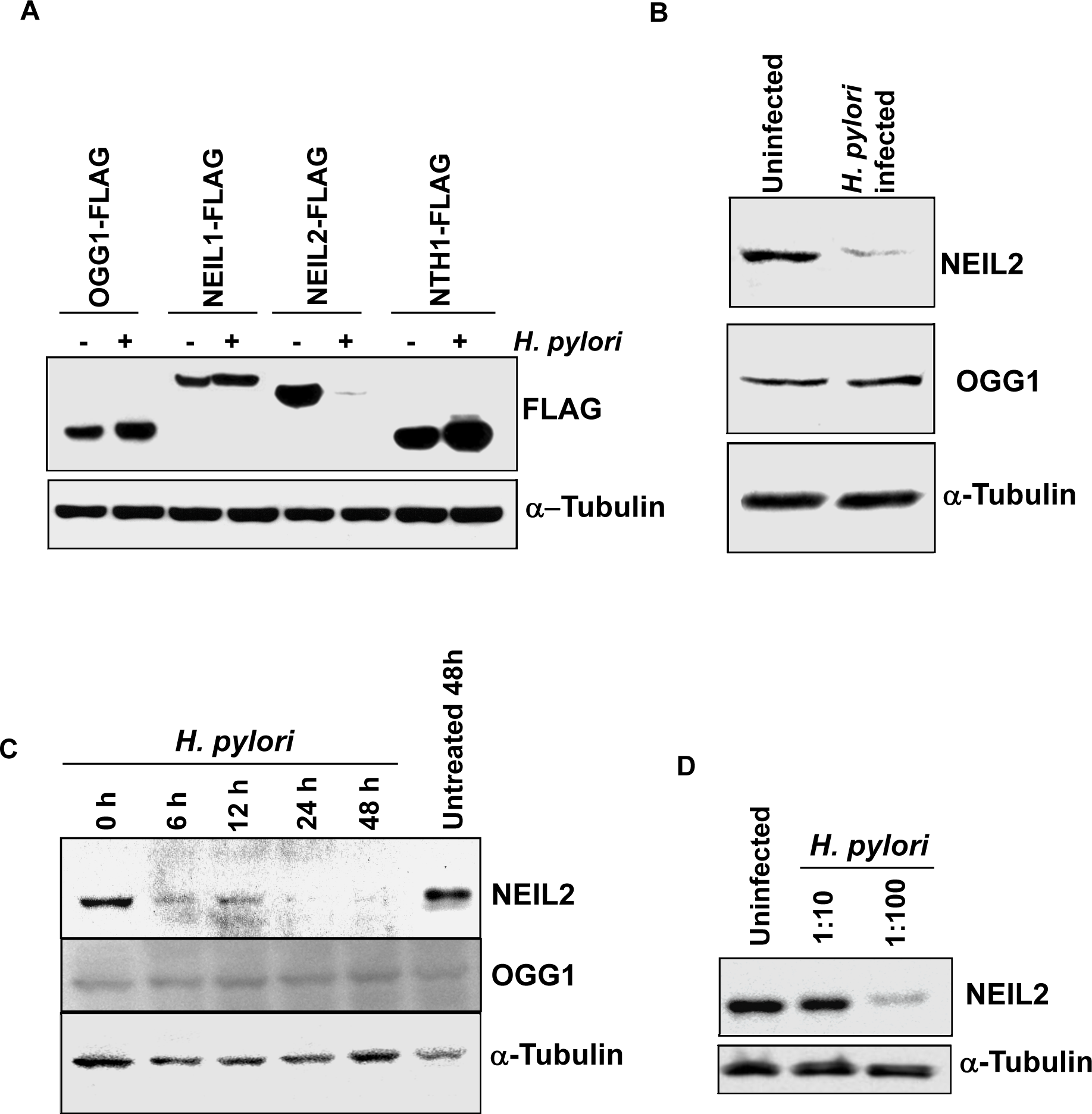
The down-regulation of NEIL2 in gastric epithelial cells after *H. pylori* infection. (A) FLAG-NEIL2, FLAG-NEIL1, FLAG-OGG1, and FLAG-NTHI were transfected in AGS cells, followed by infection with *H. pylori*. FLAG expression was determined by Western Blot (WB), with α-Tubulin used as a loading control. (B) Endogenous levels of NEIL-2 and OGG1 proteins were determined by WB in uninfected and *H. pylori*-infected AGS cells. α-Tubulin was used as a loading control. (C) Time-dependent changes in the NEIL2 expression was determined in AGS cells after infection with *H. pylori* for the indicated time points. (D) The effect of *H. pylori* MOI on NEIL2 expression was evaluated in AGS cell by using *H pylori* MOI 10:1 and 100.1.

### H. pylori infection in primary and transformed gastric epithelial cell lines down regulates NEIL2’s transcript level

To understand the impact of *H. pylori* infection on NEIL2 expression in primary and transformed gastric epithelial cell lines (Fig. 2A), we used AGS, NCI-N87, and HS738. Fig. 2B demonstrates the time-dependent changes of NEIL2 transcript level after *H. pylori* infection in AGS cells for 0, 1, 3, 6, and 12 h. The NEIL2 mRNA expression was increased after 1 h of infection, followed by a significant decrease at the transcript level in a time-dependent manner (Fig. 2B). Similarly, the level of NEIL2 transcript was also down-regulated in Hs738, the primary gastric epithelial cell line (Fig. 2C), and in NCI-N87, the transformed epithelial cell line, after 12 h of *H. pylori* infection (Fig. 2D). Next, we evaluated the physiological relevance of this finding in human gastric biopsy specimens, either *H. pylori* positive or *H. pylori* negative. Interestingly, the *H. pylori*-infected gastric biopsy specimens had lower NEIL2 expression compared to the *H. pylori-*negative subjects (Fig. 2E), indicating the link of *H. pylori* infection with the level of NEIL2 in the stomach.

**Figure 2.**
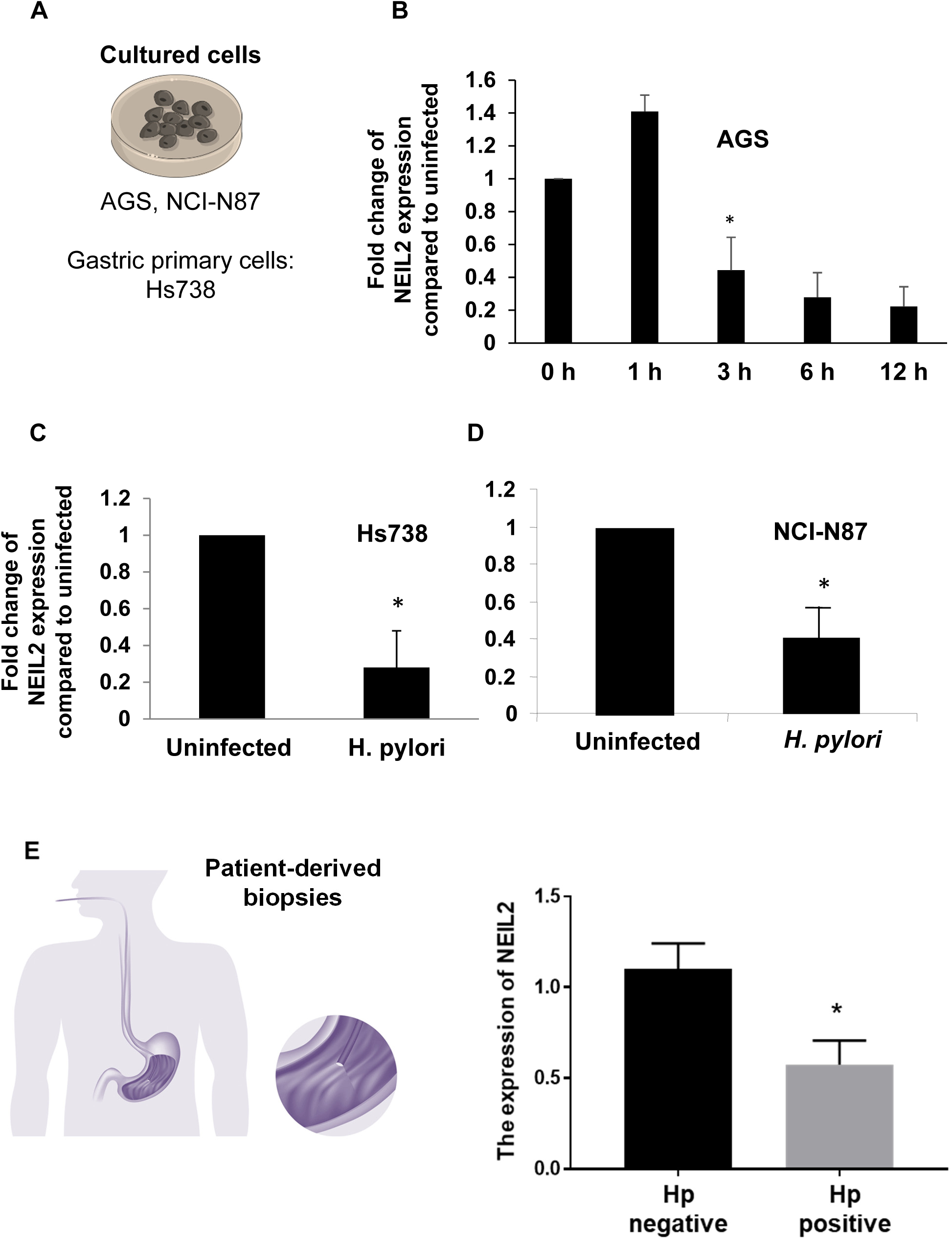
The down-regulation of NEIL2 transcript in primary and transformed gastric epithelial cells after *H. pylori* infection. (A) The schematic shows the infection of the primary and transformed cells with *H. pylori.* (B) NEIL2 transcript expression was determined by qRT-PCR (qPCR) in AGS cells after *H. pylori* infection at different time points. (C-D) NEIL2 transcript expression was tested in primary Hs738 cells (C) and NCI-N87 cells (D) after *H. pylori* infection. (E) NEIL2 expression was detected by -qPCR in *H. pylori-*positive and -negative human gastric biopsy specimens (6 negative and 4 positive). Data was generated from the mean ± SD of three independent experiments where * indicates p value<0.05.

### The down-regulation of NEIL2 is H. pylori-specific and independent of Cag pathogenicity island

To investigate whether the expression of NEIL2 was *H. pylori* infection-specific, the NEIL2 expression level was measured in AGS cells infected with either *H. pylori* or *Campylobacter jejuni (C. jejuni)*, a gastrointestinal pathogen similar to *H. pylori* (34). We found that NEIL2 expression was down-regulated with *H. pylori* infection but not with *C. jejuni* infection (Fig. 3A). To investigate if the NEIL2 expression depended on specific genes or the virulence factor of *H. pylori*, we selected two mutant *H. pylori* strains: 1) AhpC [alkyl hydroperoxide reductase, an enzyme that is crucial for oxidative stress resistance and the survival of the bacterium in the host and that generates higher amounts of 8-oxo-guanine associated with their DNA (35)]; and 2) NapA [proteins known to regulate oxidative stress and ROS generation (36)]. We also tested *H. pylori* mutant strains for Cag or CagA (*H. pylori* virulence factors) and NEIL2 expression. and compared to WT *H. pylori.* We found that NEIL2 expression was down-regulated in all of the mutants strains to a similar level to that of WT *H. pylori* (Fig. 3B-3D), indicating that the down-regulation of NEIL2 was independent of bacterial virulence factors. To understand the effect of bacterial-secreted factors, bacterial supernatant and pellet were added to the cells, and the expression was compared with the whole bacteria. The decrease in NEIL2 expression in epithelial cells was mediated only by bacterial pellet (Fig 3E, second to last lane), and boiling the pellet abolished the effect (Fig 3E, last lane), indicating that the bacterial factor responsible for NEIL2 down-regulation is heat-sensitive.

**Figure 3.**
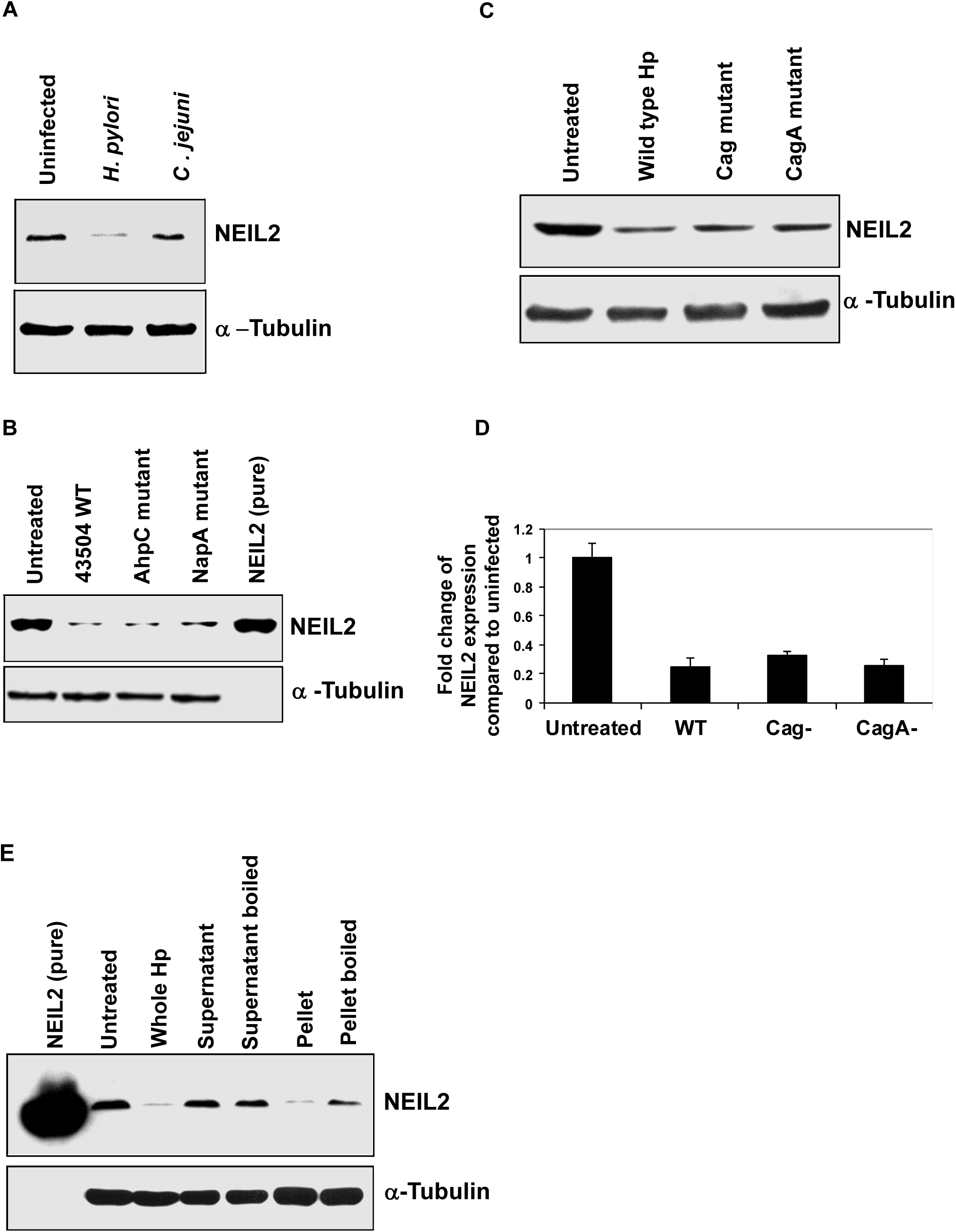
The down-regulation of NEIL2 was *H. pylori*-specific and did not depend on Cag pathogenicity island. NEIL2 expression was determined by WB in AGS cells after (A) *H. pylori* and *C. jejuni* infection, (B) 43504 WT, AhpC, and NapA mutant strains infection, and (C) WT, Cag, and CagA mutant strains infection. (D) Fold change of NEIL2 transcript expression in WT, Cag, and CagA mutant infected AGS cells was determined by qPCR. (E) NEIL2 expression was determined by WB after treatment of AGS cells with whole *H. pylori*, supernatant from infected cells, boiled supernatant, bacterial pellet, and boiled bacterial pellet. In all of the Western blots (A-E), *a*-Tubulin was used as a loading control. One representative blot from 3 independent experiments is shown.

### The inflammatory responses of gastric epithelial cells following H. pylori infection is NEIL2-dependent

To understand the link between *H. pylori*-mediated inflammatory responses and the role of NEIL2 therein, we transfected AGS cells with a vector control or NEIL2-overexpressing plasmids and infected both with *H. pylori* for 12 and 24 h. The multiplex cytokine assay showed IL-8 as the prevalent pro-inflammatory cytokine after *H. pylori* infection (not shown). Therefore, IL-8 ELISA was performed with control vs. NEIL2 over-expressing AGS cells and NEIL2 over-expression resulted in lower levels of the inflammatory cytokine IL-8 (Fig. 4A). To examine the *in vivo* significance, WT and *Neil2* KO mice were infected with *H. pylori* via oral gavage (as shown in Fig. 4B). KC (mouse IL-8 analog) mRNA expression level was measured in the gastric tissue by qRT-PCR (Fig. 4C). The level of KC expression was ∼6-fold higher in gastric tissue collected from the *Neil2* KO infected mice compared to WT infected mice (Fig. 4C). To evaluate the role of NEIL2 in gastric lesions following *H. pylori* infection, we performed H&E staining of the gastric specimens. Histology score was measured on the basis of inflammation, epithelial defects, atrophy, and metaplasia (Fig. 4D). Representative images indicate the differences in the extent of infiltration of immune cells (Fig. 4E).

**Figure 4.**
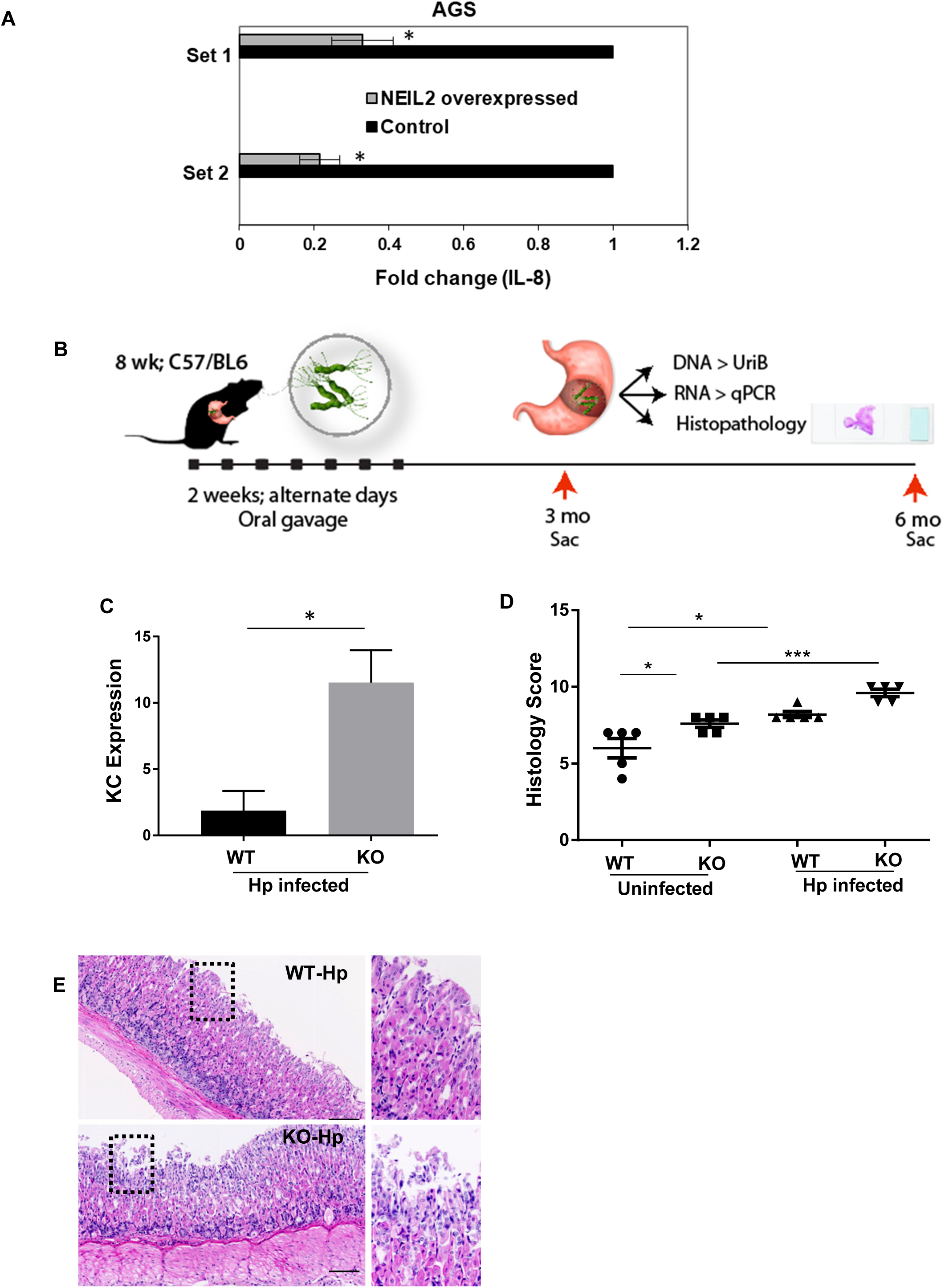
The pro-inflammatory responses in gastric epithelial cells following *H. pylori* infection depends on NEIL2. (A) The level of IL-8 (KC) was determined by ELISA in supernatant from AGS cells transfected with either vector control or full-length NEIL2 followed by infection with *H. pylori* for 12 or 24 h. (B) Schematics showing the planning of animal experiments with C57-BL/6 to isolate DNA, RNA, and histopathology. (C) IL-8 (KC) mRNA expression was determined by qPCR in gastric tissues of *H. pylori-*infected WT mice and *H. pylori*-infected *NEIL2* KO mice. (D) Histology scores of WT mice and *NEIL2* KO mice before and after *H. pylori* infection. * indicates p value<0.05, *** indicates p value<0.0005 (E) Representative images from the uninfected and infected groups.

### Enteroid-derived monolayers from NEIL2 KO mice stomach showed higher inflammation after H. pylori infection and increased DNA damage

To further confirm our results, we established EDMs from gastric spheroids (Fig. S1A) of WT and *Neil2* KO mice, then the monolayers (Fig. S1B) were infected with *H. pylori,* and KC mRNA expression was tested in both cell types (Fig. 5A). Similar to the mice data, we found that KC expression was indeed higher in *Neil2*-KO EDMs compared to WT EDMs (Fig. 5B), suggesting that NEIL2 plays an important role in mediating the inflammatory response against *H. pylori* infection. DNA damage was assessed by LA-qPCR in two representative *Polβ* and *β-globin* genes in the genomic DNA extracted from mice stomach epithelial enteroids upon *H. pylori* infection (Fig. 5C). We observed greater oxidative damage accumulation in both the *Polβ* and *β-globin* genes, in NEIL2 KO enteroids compared to WT enteroids, and also found that the damage was proportional to the duration of *H. pylori* infection.

**Figure 5.**
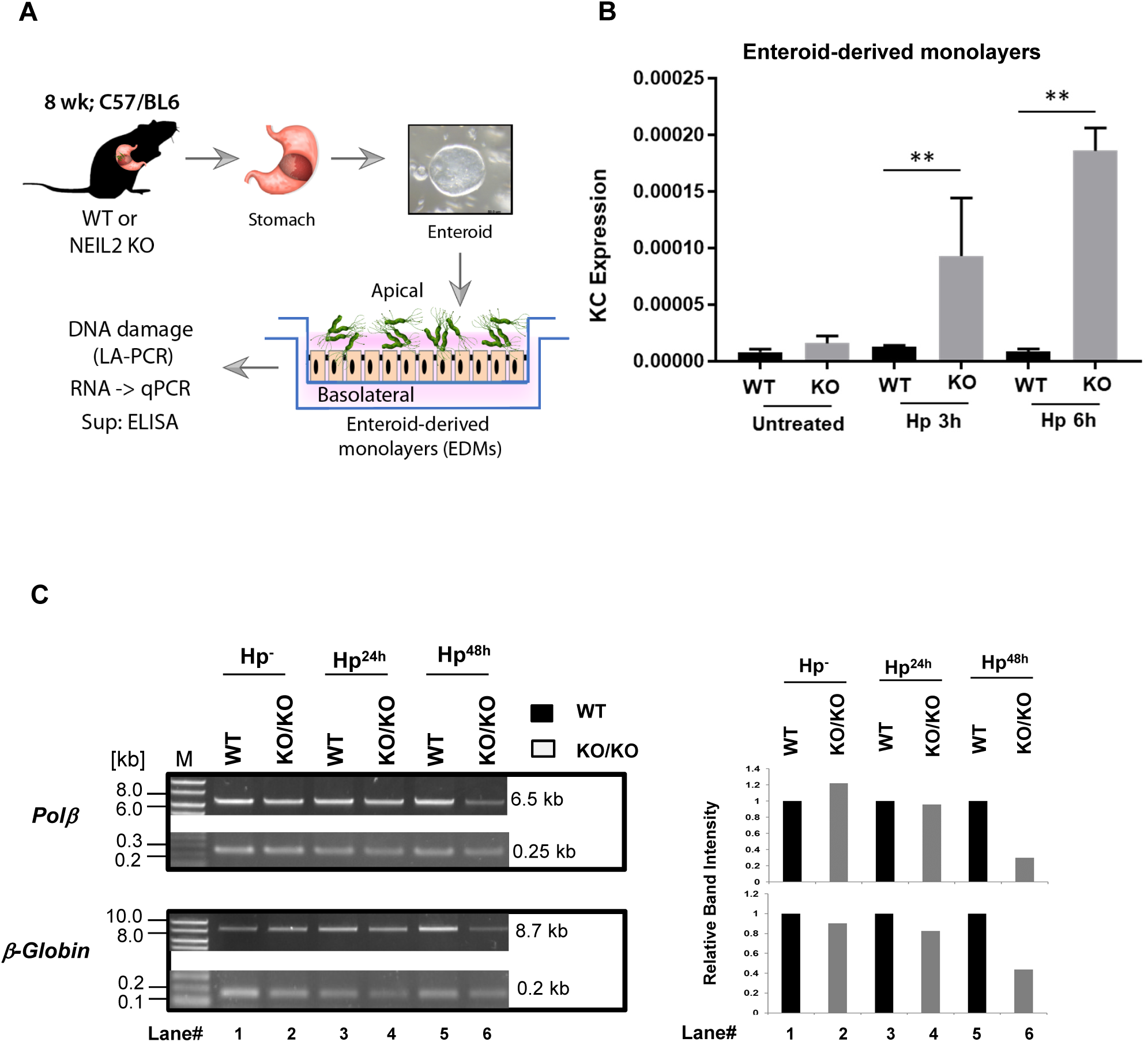
Evaluation of *H. pylori* infection-induced oxidized DNA bases in WT vs *Neil2* KO mice stomach epithelial enteroids by LA-qPCR. (A) The schematic shows the plan for enteroids isolated from WT or *Neil2* KO mice to proceed further for functional assays. (B) IL-8 (KC) expression was determined in the uninfected and infected WT and *Neil2*-KO murine enteroid-derived monolayers by Real-Time RT-PCR. ** indicates p value<0.005 (C) Stomach epithelial enteroids of WT and *Neil2*-KO mice were either mock infected or infected with *H. pylori* for 24h, 48h or 72h, and harvested for genomic DNA isolation. LA-qPCR was performed to evaluate the level of oxidized DNA base lesions after *H. pylori* infection. Representative gels show the amplification of each long fragment (∼7-8 kB) **(upper panel)** normalized to that of a short fragment (∼250 bp) **(lower panel)** of the corresponding (*Pol β* and *β-Globin*) genes. Histogram represents the DNA base damage after densitometry of the representative image from 4 different sets of WT vs. *Neil2*-KO mice. The band intensity in the WT mice arbitrarily set as unity (n=1). The error bars represent the standard deviation of the mean band intensity.

### Elevated NEIL2 gene expression in gastric cancers is correlated to higher probability of disease-free survival and NEIL2 is lowered in H. pylori-positive atrophic gastritis

NEIL2 mRNA expression levels in gastric cancers were analyzed using publicly available RNA sequencing datasets at Oncomine.org. NEIL2 mRNA expression levels were higher in both intestinal and diffuse type gastric adenocarcinomas when compared to the non-cancerous gastric mucosa (Fig. 6A, B). However, analysis of the data from the disease-free survival among 631 patients diagnosed with gastric cancers, of varying clinical stages (37) across stage I alone, stage II alone, and all stages of gastric cancers, showed that lower NEIL2 expression correlated with lower probability of disease-free survival (Fig. 6C-E).

**Figure 6.**
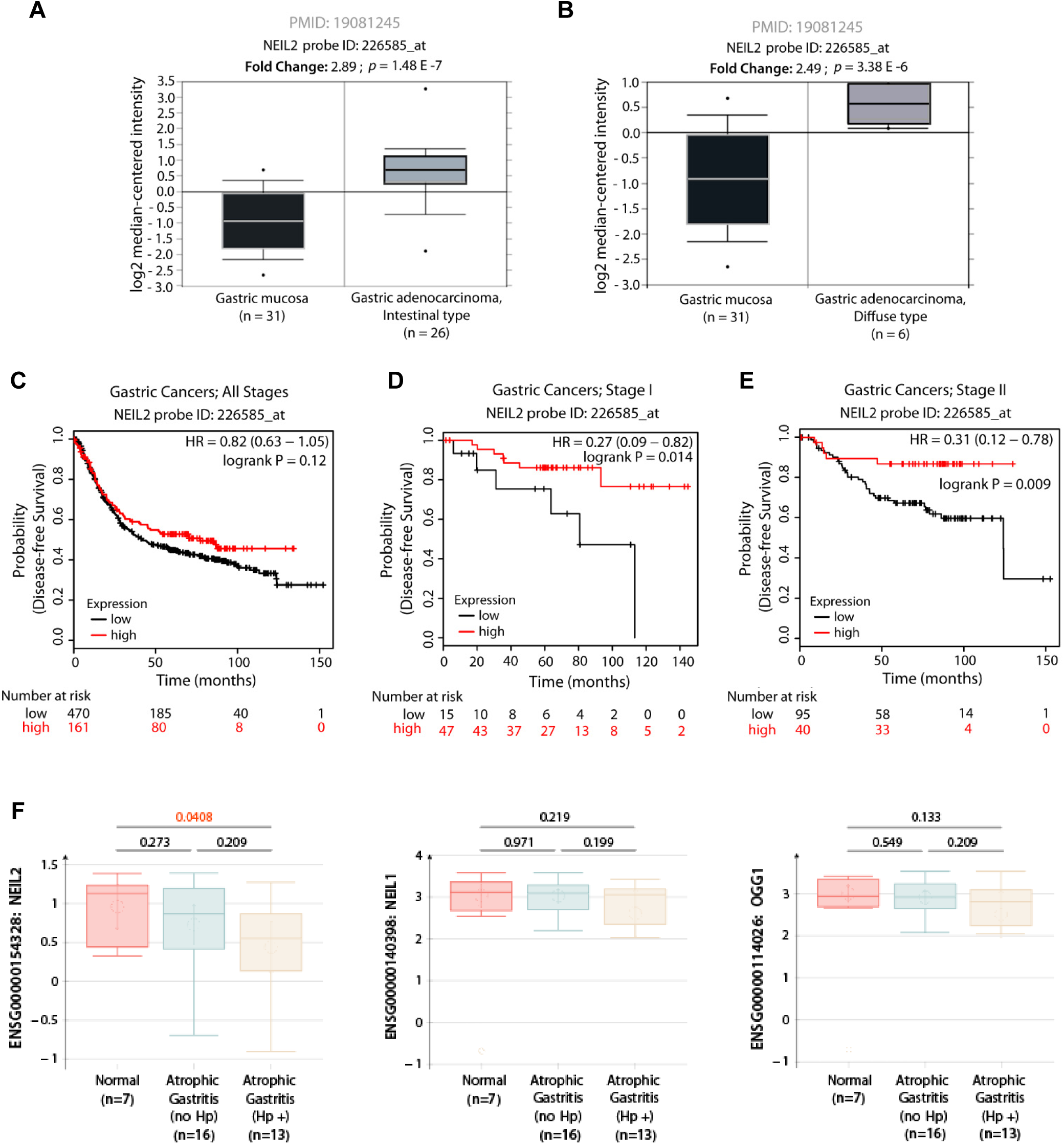
NEIL2 level is specifically down-regulated in H. pylori-positive atrophic gastritis, and low levels of expression in early gastric cancers carries a poor prognosis. A-B. NEIL2 mRNA expression is elevated in both intestinal (A) and diffuse (B) subtypes of gastric cancers. Expression levels of NEIL2 mRNA in normal vs. cancer mRNA was analyzed in publicly available RNA Seq datasets (52) using Oncomine.org. (C-E) The Kaplan-Meier plotter [K-M plotter; kmplot.com] program was used to analyze disease-free survival among 631 patients diagnosed with gastric cancers, of varying clinical stages (37) stratified into high (red) vs. low (black) NEIL2 expression. All stages, C; Stage I alone, D; Stage II alone, E. (F) The level of NEIL2, NEIL1, and OGG1 were analyzed in the publicly available dataset with GDS5338 (38) from the biopsy specimens of stomach specimens of *H. pylori*-positive and -negative subjects.

### The analysis of publicly available data identified that NEIL2 is specifically decreased with gastric cancer progression compared to other repair proteins

The publicly available dataset with record no GDS5338 (38) from the biopsy specimens of stomach from *H. pylori*-positive and -negative human subjects were analyzed, and the level of NEIL2 was found to be significantly decreased in *H. pylori*-positive gastric atrophy compared to the negative specimens. Interestingly, the down-regulation of BER proteins in gastric atrophy was prominent only for NEIL2 but not for the other two DNA glycosylases, NEIL1 and OGG1 (Fig. 6F).

Computational analysis of gene expression profiles of publicly available datasets (GSE78523) (39) showed specifically the level of *Neil2* is reduced compared to other base-excision and mismatch repair genes in gastric specimens collected from healthy subjects and patients with metaplasia (Fig. 7A). The reduction of NEIL2 correlates with the progression of gastric cancers and suggests that the reduced level of *Neil2* can accelerate mutation accumulation initiating cancer progression.

**Figure 7.**
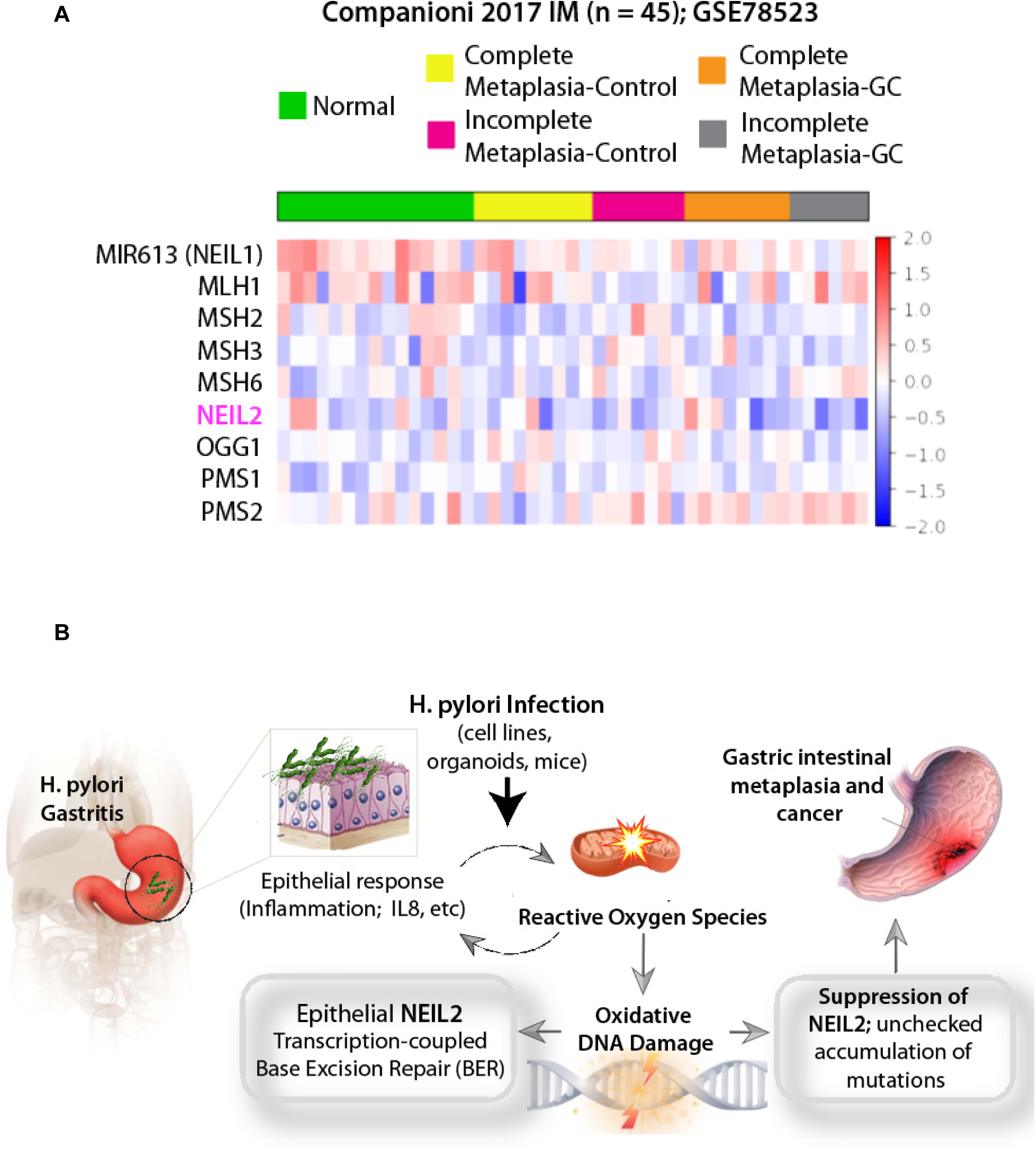
The heat map and the schematic summarizes the role of NEIL2 in the initiation of gastric cancer. (A) Heatmap is generated after analyzing the expression profile of base-excision and mismatch repair genes in the microarray of the publicly available dataset (GSE78523) (39). The microarray was performed with the RNA extracted from FFPE cuts of gastric biopsies from a total of 45 human subjects from different disease group: healthy, incomplete metaplasia and complete metaplasia. Incomplete metaplasia has higher progression rate to gastric cancer (GC) compared to the complete metaplasia. The level of NEIL2 is significantly reduced compared to all the genes analyzed in the heatmap. (B) *H. pylori* infection has serious negative effects on the gastric mucosa due to chronic mucosal inflammation leading to DNA damage and down-regulation of the base-excision repair protein NEIL2. NEIL2, which actively repairs the damage that occurs during transcription, has impaired repair activity following *H. pylori* infection and therefore accumulates mutations that can lead to gastric cancer.

## Discussion

Chronic infections and infection-induced pro-inflammatory cytokines can lead to the production of ROS and reactive nitrogen intermediates. These reactive species can induce DNA damage and thereby propel genomic instability during tumor initiation and progression (12,40). The vicious feed-forward cycle of inflammation and oxidative stress continuously targets cellular macromolecules, including the gDNA, inducing mostly oxidized base damage. These oxidized DNA base lesions are primarily repaired via the BER pathways (41,42). Defects in BER will adversely affect cellular function, accumulate DNA damage, and may trigger the development of cancers. Many carcinopathogens utilize this paradigm to induce inflammation induced genomic instability (43,44), via mechanisms that remain poorly understood. The major discovery we report here is a mechanism by which the carcinopathogen *H. pylori* derails a host defense mechanism that is geared to protect cells from oxidized DNA base lesions by repairing them.

*H. pylori* is a major human pathogen causing chronic, progressive gastric mucosal damage and is linked to gastric atrophy (AG) (45). AG is linked to an increased risk for intestinal-type adenocarcinoma and type 1 gastric carcinoids, specifically in the presence of extensive intestinal metaplasia (46). Studies have shown that eradication of the bacteria can reduce the risk of cancer development (5,47). Since *H. pylori* infection generally develops into a chronic infection, the accompanying persistent inflammatory response, ROS production, and accumulation of DNA damage could be a possible mechanism for the development of gastric cancer (48,49). In a normally functioning cell, the oxidative damage to the DNA induced by *H. pylori* infection is repaired by the BER pathway. Previous report showed that the lowered level of BER protein APE1 accumulates damage in mitochondrial DNA and the precise coordination of BER proteins in the repair of DNA damage is critical for the maintenance of genomic stability following *H. pylori* infection. (50,51). There is no study to date identified the link between BER proteins with accumulated mutation and their involvement in infection-mediated cancer. We here studied the BER protein NEIL2, as *Neil2*-null mice show the accumulation of oxidative genomic damage, mostly in the transcribed regions, and these mice are more responsive to inflammatory agents than WT mice (24). Interestingly, in gastric epithelial cells, the *H. pylori*-induced accumulation of unique DNA damage pattern occurs preferentially in transcribed regions (6).

In the current study to find the link between *H. pylori* infection and DNA glycosylases, we found that NEIL2 was the most significantly down-regulated DNA glycosylase in the human gastric epithelial cells infected with *H. pylori*. *H. pylori* down-regulated NEIL2 expression in a time- and dose-dependent manner, but was Cag independent; consequently, DNA damage was accumulated that could lead to gastric cancer. In our study, we showed the link between NEIL-2 expression and inflammatory response (IL-8, KC); overexpression of NEIL2 led to a lower inflammatory response, while *Neil2* KO mice had a greater inflammatory response. Therefore, we hypothesized that NEIL2 expression is linked to both indirect DNA damage caused by unchecked inflammatory response as well as to a direct build-up of DNA damage by inhibition of BER. This implicates the possible involvement of NEIL2 in gastric carcinogenesis after *H. pylori* infection, and it could be a potential biomarker for gastric cancer progression and prevention. From a database search, we found that NEIL2 is expressed at higher levels in various gastric adenocarcinomas compared to normal gastric tissue. However, higher NEIL2 expression within each stage of the cancer led to higher probability of survival compared to patients with lower NEIL2 expression (52). In addition to being a potential biomarker for risk and the diagnosis of carcinogenesis, NEIL2 expression may act as a prognostic biomarker to aid in determining the course of the disease. In future studies, it will be important to determine NEIL2 expression after *H. pylori* infection and correlate expression levels to gastric carcinogenesis, to identify a more significant and clinically relevant link between *H. pylori* infection, the NEIL2 protein, and gastric carcinogenesis.

Besides NEIL2 down-regulation, we found *H. pylori* infection increased the inflammatory response (IL-8 production) *in vitro* (AGS cell, Hs738, and enteroid-derived monolayers) and *in vivo* (human biopsy samples and mice). Since inflammation could be a risk factor for cancer development (44,53), we hypothesized that *H. pylori* could promote cancer progression by increasing inflammatory signaling. To link IL-8 and NEIL-2, NEIL2 knock-out (KO) mice and WT mice were tested for the inflammatory cytokine KC (CXCL1/IL-8) upon infection with *H. pylori*; we found that NEIL2 KO mice had a greater inflammatory response than WT. We confirmed our finding by infecting EDM prepared from WT and *Neil2* KO mice with *H. pylori*. Since IL-8 levels decreased in the NEIL2 overexpressed cells and increased in *Neil2* KO mice or enteroid-derived cells, confirming the regulatory role of NEIL2 on inflammation, these findings (summarized in Fig. 7B) may lead to a better understanding of infection and inflammation-mediated cancers and consequently early detection of gastric cancer and a potential cancer therapeutic. BER proteins such as NEIL2 may be utilized as important cancer drug therapeutics once the specific mechanisms through which changes in their expression levels may induce cancer formation is characterized more thoroughly. Further understanding of the exact link between *H. pylori-*mediated inflammation and gastric carcinoma through the effect of infection on the BER pathway will allow us to develop novel cancer treatments and therapies.

The findings here will pave the way for future studies in understanding the mechanism of infection-linked carcinogenesis and open avenues for studying the link between infection-mediated inflammation and cancer. We provide a perspective on DNA repair inhibition and inflammatory response and propose that the two are regulated by NEIL2 and exacerbate one another to foster tumorigenic DNA damage. More research is needed to investigate the link between DNA repair and infection-associated inflammatory response in oncogenesis. Finally, the concepts discussed here can be studied in and applied to the many other infection-associated cancers.

## Materials and Methods

### Spheroid Isolation and Maintenance

A spheroid of gastric stem cells were isolated from wild type (WT) and *Neil2* KO mice, as described previously (54,55). The pylorus section of the stomach was removed, washed with PBS and then incubated with collagenase type I solution containing gentamicin (50 ug/ml) at 37°C for 30 min with mixing at 10-min intervals. The cells were filtered through a 70-µm cell strainer, and the density and viability of the gastric stem cells were checked using trypan blue and hemocytometer. The cells were suspended in Matrigel, and 50% conditioned medium (CM) was added to each well (54). Spheroid cultures were passaged every three to four days using 0.25% trypsin in PBS-EDTA. Enteroid-derived Monolayer (EDM) was prepared as previously described (20,56). The spheroids were briefly trypsinized, and the isolated cells were filtered, counted, suspended in 5% CM and pelleted with Matrigel (1:30 dilution) in the apical part of the transwell at the density of 5×10^5^ cell/well. The 5% CM was added to the base of the transwell.

### Cell Line Maintenance

AGS cells were cultured in RPMI 1640 medium containing 10% fetal bovine serum (FBS), 1 mM glutamine, penicillin (1 U/mL), and streptomycin (100 µg/mL) and kept in an incubator at 37°C and 5% CO_2_.

### H. pylori Infection

*H. pylori* strain (ATCC-26695) was incubated with the Brucella Broth supplemented with 10% FBS one day before the infection and left on a shaker at 37°C in 10% CO_2_. EDM and AGS-cells were infected with the bacteria at a multiplicity of infection (MOI) of 100:1.

### RNA Isolation and RT-PCR

RNA was isolated from uninfected and *H. pylori*-infected cells using the RNeasy Micro Kit (Qiagen, Valencia, CA) according to the manufacturer’s instructions. cDNA was synthesized using the superscript kit (Invitrogen). Quantitative PCR (qPCR) was carried out using Syber Green master mix for target genes and normalized to the housekeeping genes GAPDH or B-actin using the ∆∆C_t_ method [30]. Primers were designed with Roche Universal Probe Library Assay Design software.

### Western Blot

Cell lysates, collected from uninfected and *H. pylori*-infected monolayers and cell lines, were tested for NEIL2 protein expression using purified anti-NEIL-2 antibody (dilution 1:500) as described previously (57).

### ELISA

Supernatants, collected from apical and basolateral parts of uninfected and *H. pylori*-infected monolayers on transwells, were tested for IL-8 using the mouse CXCL1/KC DuoSet ELISA kit (R&D Systems) according to the manufacturer’s instructions. The level of IL-8 was compared in *H. pylori*-infected EDMs versus the uninfected EDMs with the same cell number.

### Infection of Neil2-KO mice with H. pylori

Mice used in the study were WT and *Neil2* KO C57BL/6 mice and were generated and characterized as previously described (24). The maintenance, breeding, and infection studies protocols were approved by the Animal Care and Ethics Committee of UTMB, Galveston, TX. Mice were inoculated with *H. pylori* strain SS1 (10^8^ bacterial counts/dose) via oral gavage for 3 days/week as described previously (58). Animals were sacrificed after 3 months and 6 months and stomach specimens were collected for RNA and DNA isolation.

### Biopsy Samples

Gastric biopsy samples were collected from gastro-esophageal-duodenoscopy patients according to Institutional Review Board approved human subject protocols. All the human studies abide by the Declaration of Helsinki principles. Histopathology and rapid urease testing were used to detect the *H. pylori* infection. The NEIL2 transcript expression was determined using qRT-PCR.

### DNA Damage Analysis

WT and *Neil2* KO mice were infected with *H. pylori,* and genomic DNA (gDNA) was isolated from stomach epithelial enteroids of uninfected control and *H. pylori*-infected mice using a genomic-tip 20/G kit (Qiagen, catalog no. 10223, with corresponding buffer sets) per the manufacturer’s protocol. This kit has the advantage of minimizing DNA oxidation during the isolation steps, and thus it can be used reliably for isolation of high molecular weight DNA with excellent template integrity for detecting endogenous DNA damage using Long Amplicon quantitative-PCR (LA-qPCR). The extracted gDNA was quantified by Pico Green (Molecular Probes) in a 96-well black-bottomed plate, and the presence of *H. pylori* was confirmed by PCR. To induce strand breaks at the sites of unrepaired oxidized DNA base lesions, 150 nanograms of gDNA were treated with 4 units of *E. coli* enzyme Fpg (New England Biolabs) for 1 hour at 37°C. The enzyme was inactivated by incubating the reaction mixture at 65°C for 15 min. Gene-specific DNA damage was measured by LA-qPCR using 15 nanograms of Fpg-treated gDNA per PCR reaction following the protocol as reported earlier by Chakraborty, A. *et. al.* (20). LA-qPCR was carried out to amplify a 6.5-kb region of *pol* ß and the 8.7-kb region of globin genes, as done previously (20). To maintain linearity, the final PCR condition was optimized at 94°C for 30 s (94°C for 30 s, 55–60°C for 30 s depending on the oligo annealing temperature, 65°C for 10 min) for 25 cycles and 65°C for 10 min. Since amplification of a small region would be independent of DNA damage, a small DNA fragment for each gene was also amplified to normalize the amplification of large fragment,s as done before (20). The amplified products were then visualized on gels and quantified with an ImageJ automated digitizing system (National Institutes of Health) based on two independent replicate PCRs. The extent of damage was calculated in terms of relative band intensity with the WT mice considered as unity.

### Histopathology of Neil2 KO mice following H. pylori infection

Gastric lesions were graded blindly on an ascending scale from 0 to 4 for inflammation, epithelial defects, atrophy, hyperplasia, pseudo-pyloric metaplasia, dysplasia, and mucous metaplasia, as defined elsewhere (59). A gastric histologic activity index was generated by combining scores for all criteria. Additionally, the tissue pieces were embedded in paraffin, sectioned at 5 µm and stained with hematoxylin and eosin for histologic evaluation using the same scoring system as previously described (60).

### Statistics

Results are expressed as mean ± SD and were compared by using two-tailed Student t test. Differences were considered significant if P values were lower than 0.05.

## Acknowledgment

This work was supported, in whole or in part, by National Institute of Health Grants: DK107585 and DK099275 (to SD); R01 NS073976 (to TH); and R01HL145477 (to TH); DoD CA150375 (to VER). We thank Sarah Toombs Smith, Ph.D., ELS, for critically editing this manuscript.

## Abbreviations

GC: Gastric cancer
EDMs: Enteroid-derived monolayers
MOI: Multiplicity of Infection
WT: Wild Type
BER: Base Excision Repair
DGs: DNA glycosylases
LA-qPCR: Long Amplicon quantitative-PCR
AG: Gastric Atrophy.

## Figure Legends

**Supplementary Figure, Figure S1.**
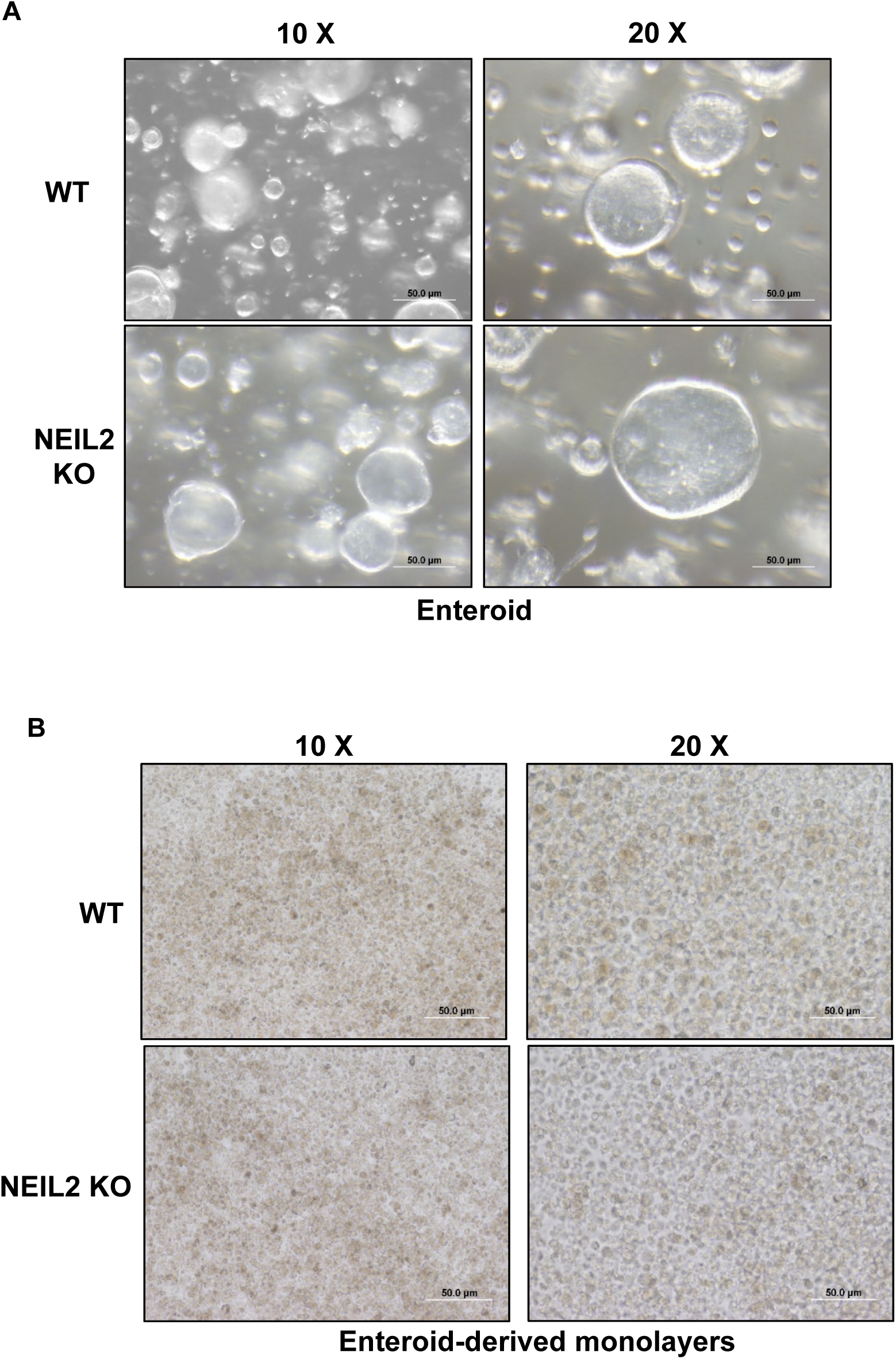
Gastric organoid, and organoid-derived monolayers. (A) Enteroids isolated from gastric specimens of WT and *Neil2* KO mice were polarized to monolayers, enteroid-derived monolayers (B). A representative picture has shown with two different magnification (10 X, 20 X).

